# More is better: A simple antibody-based strategy for recovering all major mouse brain cell types from multiplexed single-cell RNAseq samples

**DOI:** 10.1101/2024.06.11.598585

**Authors:** Federico Pratesi, Laura K. Hamilton, Jessica Avila Lopez, Anne Aumont, Mariano Avino, Nishita Singh, Ihor Arefiev, Marie A. Brunet, Karl J. L. Fernandes

**Affiliations:** Research Center on Aging, CIUSSS de l’Estrie-CHUS, Sherbrooke, Canada; Department of Medicine, Faculty of Medicine and Health Sciences, Université de Sherbrooke, Sherbrooke, Canada; Research Centre of the Université de Montréal hospital (CRCHUM), Montreal, Canada; Bio-informatic Platform, Department of Biochemistry and Functional Genomics, Faculty of Medicine and Health Sciences, Université de Sherbrooke, Sherbrooke, Canada

## Abstract

Single-cell RNA sequencing (scRNAseq) is a powerful yet costly technique for studying cellular diversity within the complexity of organs and tissues. Here, we sought to establish an effective multiplexing strategy for the adult mouse brain that could allow multiple experimental groups to be pooled into a single sample for sequencing, reducing costs, increasing data yield, and eliminating batch effects. We first describe an optimized cold temperature single-cell dissociation protocol that permits isolation of a high yield and viability of brain cells from the adult mouse. Cells isolated using this protocol were then screened by flow cytometry using a panel of antibodies, allowing identification of a single antibody, anti-Thy1.2, that can tag the vast majority of isolated mouse brain cells. We then used this primary antibody against a “universal” neural target, together with secondary antibodies carrying sample-specific oligonucleotides and the BD Rhapsody single-cell system and show that multiple adult mouse brain samples can be pooled into a single multiplexed run for scRNAseq. Bioinformatic analyses enable efficient demultiplexing of the sequenced pooled brain sample, with high tagging efficiency and precise annotation and clustering of brain cell populations. The efficiency and flexibility of the cell dissociation protocol and the two-step multiplexing strategy simplifies experimental design, optimizes reagent usage, eliminates sequencing batch effects and reduces overall experimental costs.

## INTRODUCTION

Single-cell RNA sequencing (scRNAseq) is a technologically advanced technique that is now established as a potent strategy for investigating cellular diversity and tissue lineage development^1–3^. It is particularly useful for studying highly heterogeneous tissues such as the brain, which is composed of multiple types of neural lineage cells (neurons, astrocytes, oligodendrocytes, ependymal cells, neural precursors) as well as resident immune cells (microglia), vasculature associated cells (endothelial cells, pericytes, smooth muscle cells) and others^4–6^. Multiple systems for single cell isolation and manipulation have been developed to obtain single cells for RNA extraction, transcriptome preparation and sequencing^3,7,8^. These include platforms that are primarily microfluidics-based, such as 10x Genomics Chromium^9^, or microwell-based, namely the BD Rhapsody^10^. A comparative study between the 2 technologies has been performed by Salcher et al. ^11^. More companies are joining the market including HoneyComb HIVE^12^, Parse Biosciences^13^ and PIP-seq (commercialized by Fluent BioSciences)^14^, with interesting future perspectives for technological advances in this field.

Advances in the cell loading capacity of the latest versions of single-cell platforms have now made it feasible to “multiplex” several experimental groups into a single sample for scRNAseq, thus eliminating batch effects during sequencing and substantially reducing overall sequencing costs. During scRNAseq, each cell within an experimental sample is individually barcoded, allowing each sequenced cDNA fragment from that sample to be ascribed to a specific cell. Multiplexing of experimental samples requires adding a second, “sample tag” to each cell that will further identify from which experimental group it is originated ^15^. Besides directly reducing costs and sequencing variability, multiplexing has the added benefit of revealing the frequency of multiplets (presence of two sample tags) to help optimize cell loading density. Two main multiplexing methods are currently used, antibody-based and lipid-based^16^.

In the present study, we describe a straightforward, two-step, antibody-based approach for multiplexing samples specifically from the adult mouse brain. Finding multiplexing methods that can tag all cell types is particularly challenging for complex tissues that possess a wide diversity of cell types and lineages, such as the brain. The diversity of cell types and lineages in the adult brain poses a particular challenge for antibody-based multiplexing, with most antibodies targeting antigens that have cell- or lineage-specific expression patterns. Here, we show that a primary antibody against the glycoprotein Thy1.2 (also known as CD90.2) labels most neural and non-neural cells dissociated from the adult mouse brain. When combined with sample-specific oligonucleotide-labelled secondary antibodies (the BD FLEX-Single-cell Multiplexing Kit) and the microwell-based BD Rhapsody single-cell system, this enables a multiplexed sample to be effectively demultiplexed following sequencing and the majority of brain cell types to be recovered for transcriptomic analysis.

## RESULTS

### Validation of a tissue dissociation protocol for high cell viability and yield from micro dissected regions of the adult mouse brain

Cell dissociation is a critical initial step in single-cell transcriptomic experiments^17,18^. For studies on the adult brain, this step is particularly important given the highly physically interconnected nature of the cells within nervous tissue^19,20^. To maximize the quality of cells dissociated from adult mouse brain, we incorporated modifications aimed at limiting mechanical, temperature, and time-associated stresses (summarized in Figure 1a). These modifications included adding transcription inhibitors during brain dissection and dissociation, performing enzyme incubations at 4°C rather than at 37°C, reducing washing and centrifugation steps to a bare minimum to decrease processing time and stresses, and incorporating a dead cell removal step (Miltenyi columns-based kit) to enhance the proportion of viable cells. Transcription inhibitors and cold temperatures were used to minimize *ex vivo* activation of cells, as previously reported^21–23^. We also eliminated commonly used isolation and purification steps like FACS (due to its microfluidic stresses and added processing time)^24^ and Percoll gradient centrifugation (due to significant declines in cell yield). We took advantage of the BD Rhapsody scanner single-cell platform to visualize and quantify numbers of live cells (Calcein AM+) and dead cells (Draq7+), allowing immediate optimization of these steps. Figure 1 provides an overview of the dissociation protocol and quantification of cell viability and yield using the BD Rhapsody.

**Figure 1.** Validation of tissue dissociation protocol for adult mouse brain. a) Graphical abstract of the optimized dissociation protocol. This includes perfusion and brain dissection (steps 1 and 2), microdissection of the regions of interest (step 2), dissection to single cell suspension (steps 3 and 4), dead cell removal using magnetic columns (step 5) and cell counting and sample preparation (step 6). b) Calcein AM (live cells) and Draq7 (dead cells) visualized with the Rhapsody scanner. c) Charts of viability and cell concentration of samples from 10 tests in 3 brain regions.

To validate the cell dissociation protocol, we performed a series of experiments in which three restricted brain regions of interest, the hypothalamus, the hippocampus and the lateral ventricle wall (LVW), were microdissected from female 3xTg-AD mice and age-/sex-matched strain controls^25^. We tested the protocol using a range of ages, from 2 months old to 10 months old, not finding significant difference in the final cell yield (Figure 1c, and Figure S3). As an example, in a standard experiment dissociating only 2 brain regions, 3 experimenters were involved with one person perfusing and euthanizing the mice, one person dissecting out the brain, and one person microdissecting the hippocampus, hypothalamus, and/or LVW (see Methods). Samples were collected in 1.5mL Eppendorf tubes with each tube containing one microdissection from each of two mice (2 microdissections per tube). Animals were sacrificed sequentially, with the time from each sacrifice to the microdissections being placed on ice being 5-7 minutes per mouse and the total collection time for 8 mice being 1 hour (Figure 1a, steps 1 to 3). The collected samples were then dissociated in parallel (as detailed in the Methods), the dissociated cells from each experimental group pooled, and the viability assessed as described using the BD Rhapsody scanner, an additional 1.5 hour total (Figure 1a, steps 4 to 6). Across 10 separate experiments and the 3 brain regions, cell viability ranged from 87% to 97.2% (Figure 1c). Since there was no difference in cell yield between 3xTg and control mice, data from the two lines of mice are combined in Figure 1c. Mean cell yield from these regions was 348 072 ±68 025 cells/hypothalamus, 168 044 ±26 537 cells/ hippocampus and 228 680 ±72 324 cells/LVW. The variability of viability and concentration across these brain areas was non-significant (Figure S3).

### Identification of anti-Thy1.2 as a “universal” label for dissociated adult mouse brain cells

Two antibody-based single-step multiplexing kits are currently available from BD to allow multiple experimental groups to be pooled into a single sample for scRNAseq^10,26^. These kits use primary antibodies against either CD45 or MHC-I that are tagged with one of up to twelve oligonucleotide barcodes, theoretically allowing up to 12 samples to be multiplexed for single-cell RNAseq. However, testing of the CD45 and MHC-I kits on adult brain cells demonstrated restricted cell-type labelling patterns, with CD45 tagging only 41% of cells on average and efficiently tagging mainly microglia and immune cells, and MHC-I only tagging 67% on average and efficiently tagging mainly endothelial cells (Figure S1). We therefore asked whether we could devise a novel antibody-based multiplexing strategy that would enable a larger proportion of adult brain cell types to be recovered after demultiplexing.

To do so, we aimed to take advantage of the BD Flex-SMK kit^27^, which uses oligonucleotide-barcoded anti-PE secondary antibodies, and attempted to identify one or a combination of PE-labelled primary antibodies that would allow tagging of a larger percentage of dissociated brain cells (i.e., a 2-step multiplexing approach) (Figure 2a). We initially generated a list of potential neural cell type markers by cross-referencing the catalog of available PE-conjugated antibodies with widely recognized neural markers documented in the literature (Table S1). From this initial pool, we selected eight PE-conjugated primary antibodies that would be predicted to encompass the majority of cell populations we had identified in our prior single-cell experiments (see Table 1): ACSA-2 (astrocytes), Notch-1 (neuroblasts), CD140a (oligodendrocytes, pericytes), NRP-1 (vascular/leptomeningeal), MHC 1 (microglia, tanycytes, immune), CD24 (ependymal), CD13/CD140a (pericytes), CD90.2 (Thy-1.2) (neurons) (Table 1).

**Figure 2.**
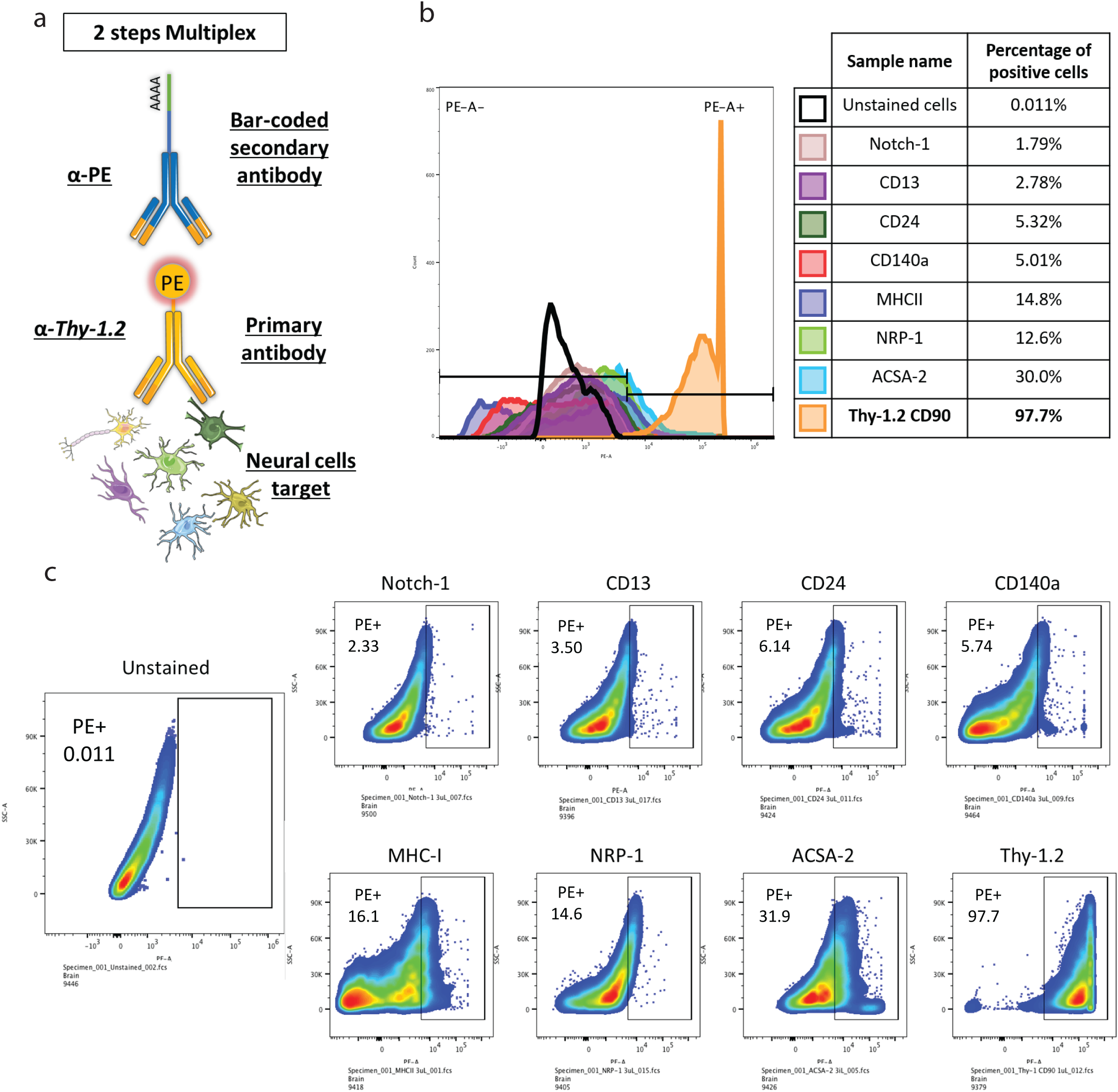
Anti-Thy1.2 labels the vast majority of dissociated adult brain cells. a) Description of double labeling strategy with Flex SMK kit. b) Modal option scales of all channels as a percentage of the maximum count. c) Gated positive populations for each tested marker using FlowJo. Note that 97.7% of all cells are Thy1.2 positive.

**Table 1:** PE-conjugated antibodies and their target cell types.

| Cell type | Target | Catalogue number | Company |
| --- | --- | --- | --- |
| Astrocyte | ACSA-2 | 130-116-141 | Miltenyi |
| Neuroblast | Notch-1 | 130608 | Biolegend |
| Oligodendrocyte | CD140a | 135905 | Biolegend |
| Vascular/Leptomeningeal | NRP-1 | 145203 | Biolegend |
| Endothelial | MHC I | 566776 | BD |
| Tanycyte |  |  |  |
| Immune (incl. Microglia) |  |  |  |
| Ependymocyte | CD24 | 567383 | BD |
| Pericyte | CD13/CD140a | 558745 | BD |
| Neuron | CD90.2 (Thy-1.2) | 553005 | BD |

Flow cytometry analysis of cells dissociated from the hippocampi of wildtype mice and labelled with the above PE-conjugated primary antibodies revealed that 7 of the 8 antibodies labelled less than 30% of hippocampal cells each. To our surprise, however, more than 97% of hippocampal cells were labelled by PE-conjugated anti-Thy-1.2 (Figure 2b-c). To validate this unexpected observation, we repeated this flow cytometry analysis on cells dissociated from additional adult brain regions and again found that anti-Thy-1.2 tagged the majority of cells from the hippocampus (97.7%), hypothalamus (95.7%) and LVW (97.7%) (Figure S2).

Thus, anti-Thy1.2 has the potential to serve as a “universal” neural marker for tagging cells dissociated from the adult mouse brain. We therefore investigated its possible usage as a multiplexing tool.

### A 2-step multiplexing strategy for the adult brain using PE-labelled anti-Thy1.2, the BD Flex-SMK kit and the BD Rhapsody single-cell platform: workflow from single-cell suspension to library preparation

To test the multiplexing potential of PE-labelled anti-Thy1.2 and the BD Flex-SMK (single-cell multiplexing kit), we microdissected the hypothalamus and LVW from 4-month-old female WT and 3xTg-AD mice, yielding 4 experimental groups for multiplexing (Figure 3a). Tissues were dissociated using the above optimized cell dissociation protocol, again yielding between 115 360 and 346 320 cells per experimental group with 83.3 – 95.4% viability (Figure 3b). Cells from each of the four experimental groups were labelled with PE-conjugated anti-Thy1.2 primary antibody, followed by one of the oligonucleotide-barcoded anti-PE antibody sample tags (Figure 3c-e and detailed in the Methods). We then pooled 15 000 cells from each group (60 000 cells total) into a single multiplexed sample for loading onto the BD Rhapsody single lane cartridge (Figure 3 f).

**Figure 3.**
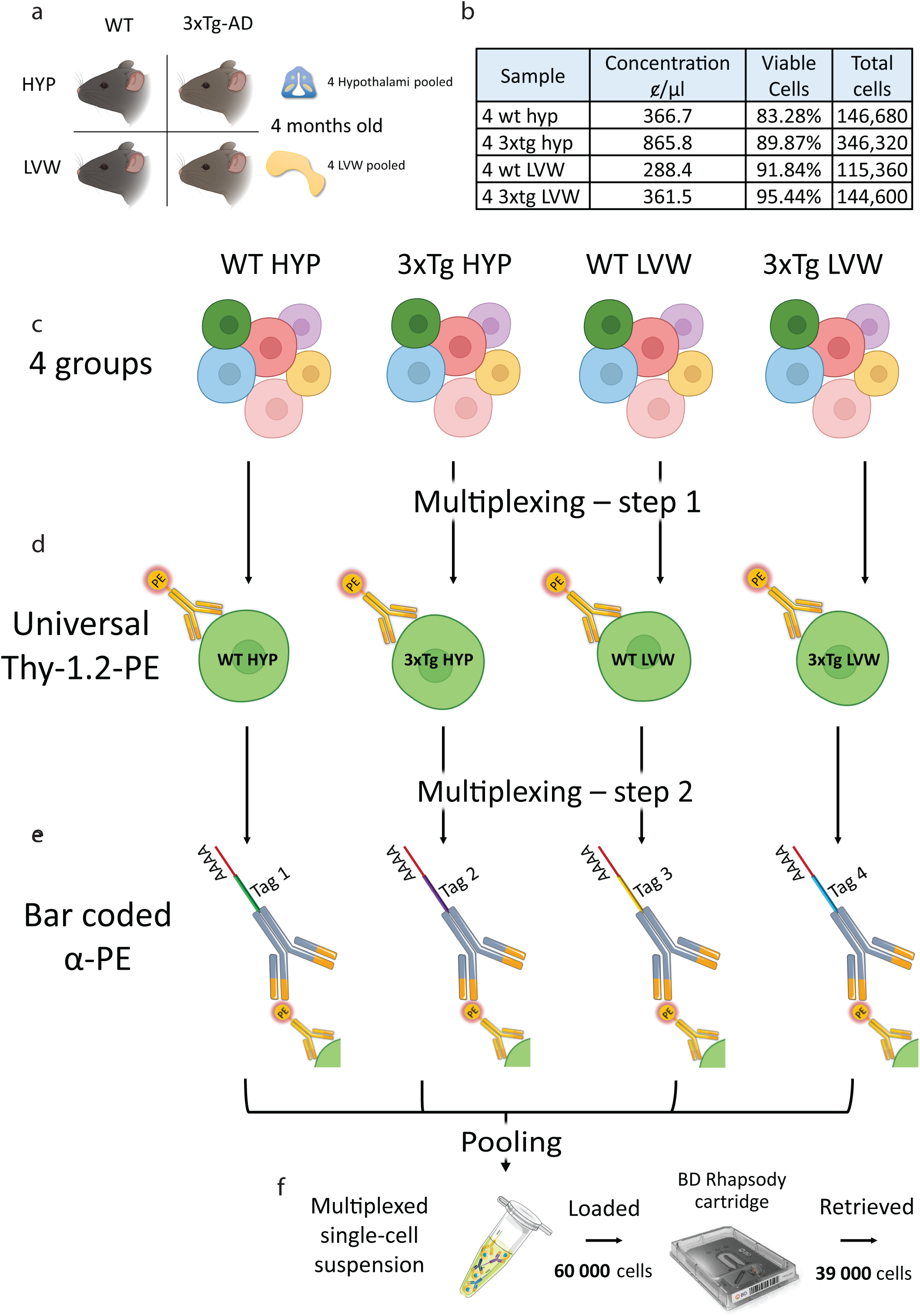
Two-step multiplexing strategy using anti-Thy1.2. a) Experimental plan showing the 4 groups for the experiment: WT and 3xTg strains and hypothalamus and LVW microdissections for each strain. b) Table showing the cell concentration and viability from the single cell suspensions. c-g) The three stages of the multiplexing procedure involve using the cell suspensions enriched for live cells (c), labelling of each sample with PE-conjugated Thy-1.2 primary antibody (d), labelling with the oligo-conjugated anti-PE secondary antibody (e), pooling the labelled cell suspension all samples into a single tube (f) and loading onto the Rhapsody cartridge. Loading 60 000 cells (15 000/sample) in the single lane cartridge allows retrieval of approximately 39 000 cells.

Results of the BD Rhapsody scanner quality control analysis are summarized in Table 2, highlighting collection of approximately 39 000 candidate single cells from the cartridge. As per the manufacturer instructions, we then generated the required Whole Transcriptome Analysis (WTA) library (i.e., cDNA amplification and preparation for transcriptomic analyses) and the corresponding sample tag library (i.e., sample tag expansion and preparation to allow demultiplexing of the experimental groups). Details of the workflow for cartridge preparation, sample loading, and barcoded bead collection are detailed in the ‘Star Methods’ section. The WTA and sample tag libraries were sequenced on a Novaseq 6000, with each cell targeted for a depth of 35 000 reads per cell, and an additional 1 200 reads per sample tag per cell. Subsequently, the alignment and initial quality control procedures were executed on the SevenBridges genomic platform (now Velsera) using a reference mouse genome (see methods).

**Table 2:** Results from BD Rhapsody scanner quality control showing vast majority of viable cells are retrieved.

|  |  |
| --- | --- |
| Number of viable cells captured in wells at cell load | 42,288 |
| Number of wells with viable cells and a bead | 39,381 |
| Cell multiplet rate | 9.00% |
| Bead retrieval efficiency | 98.80% |
| Number of retrieved beads with 1+ viable cells | 38,869 |

### Clustering, annotation and demultiplexing of four experimental groups

Following sequencing and alignment, we asked SevenBridges to identify the 35 000 best putative cells of which 4 363 were identified as multiplets (Figure S4a). After further bioinformatic quality controls for mitochondrial genes, multiplets and number of genes expressed per cell, we obtained 20 923 total cells. Utilizing Seurat dimensionality reduction and clustering workflow, and UMAP technique for the visualization of clusters, 18 cell clusters were identified (Figure 4a) and successfully annotated for cell type by comparison with previously published gene sets using a combination of automated and manual tools (see Figure 4a and table 3). The annotated cell populations include astrocytes, oligodendrocytes, microglia, endothelial cells, vascular cell, GABAergic neurons, 2 mixed populations (astrocyte_microglia and astrocyte_endothelial), and a few less represented cell types. Other publications on the same brain regions show similar numbers of clusters and annotations^10,21,28,29^.

**Figure 4.**
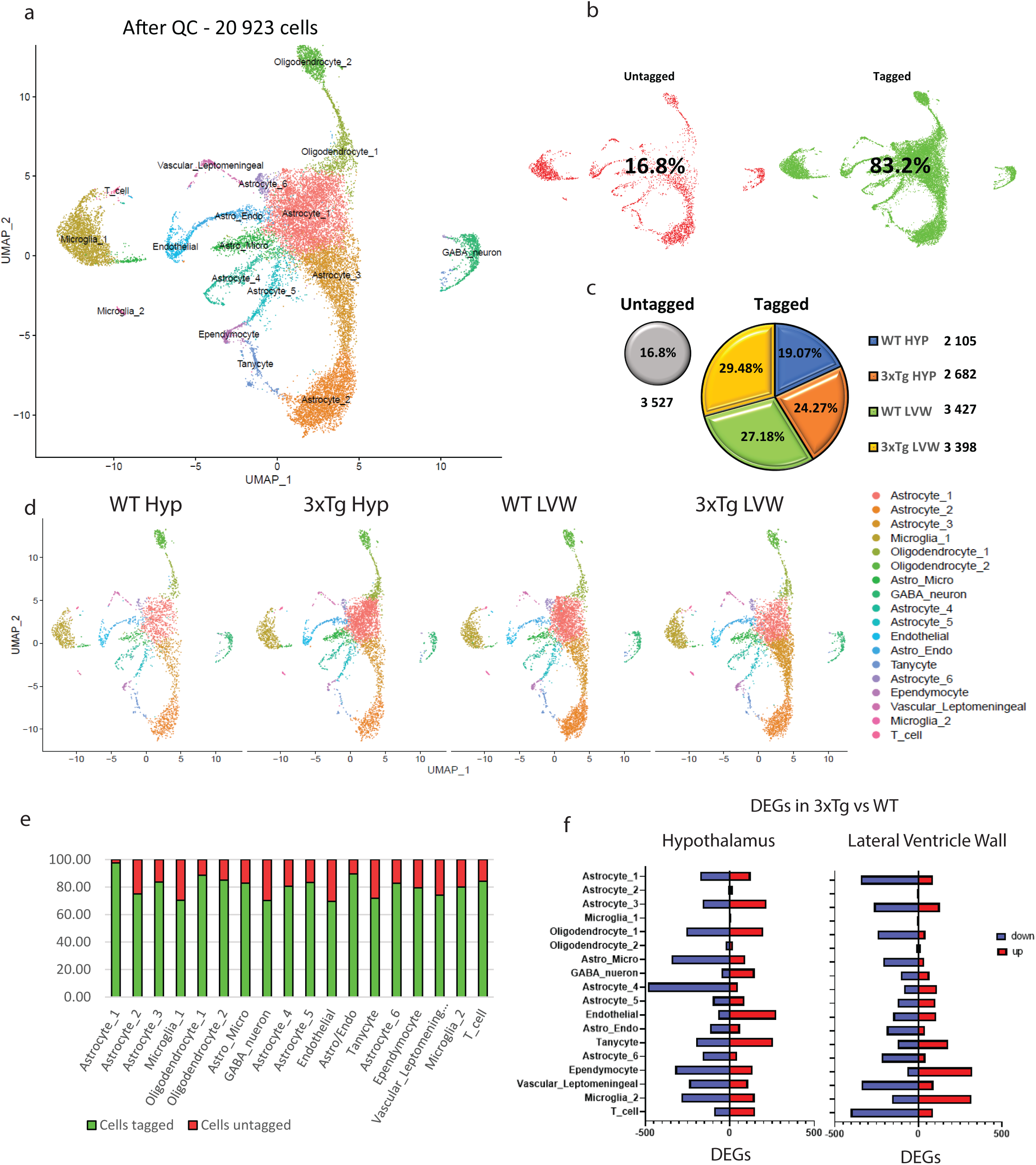
Efficient recovery of sample identity and annotation of cell types dissociated from adult mouse brain. a) UMAP of all cells after QC. b) UMAPs of untagged cells compared to tagged cells. c) Pie chart with numbers and percentages of the 4 tagged groups plus the untagged group. d) UMAPs of the 4 tagged group with corresponding clusters. e) Percentage of tagged cells per population. f) Up and down regulated differentially expressed genes per population and brain region.

**Table 3:** Master table with cell types, percentages and numbers of tagged cells per group.

|  | Tot cells | Tot cells tagged | Untagged | % tagged | WT_Hyp | WT_Hyp % on tagged | Hyp_3xTg | Hyp_3xTg % on tagged | WT_Svz | WT_Svz % on tagged | Svz_3xTg | Svz_3xTg % on tagged |
| --- | --- | --- | --- | --- | --- | --- | --- | --- | --- | --- | --- | --- |
| Astrocyte_1 | 5148 | 5020 | 128 | 97.51 | 590 | 11.75 | 1618 | 32.23 | 1674 | 33.35 | 1138 | 22.67 |
| Astrocyte_2 | 3207 | 2403 | 804 | 74.93 | 392 | 16.31 | 459 | 19.10 | 852 | 35.46 | 700 | 29.13 |
| Astrocyte_3 | 3030 | 2536 | 494 | 83.70 | 316 | 12.46 | 472 | 18.61 | 642 | 25.32 | 1106 | 43.61 |
| Microglia_1 | 2638 | 1857 | 781 | 70.39 | 382 | 20.57 | 455 | 24.50 | 487 | 26.23 | 533 | 28.70 |
| Oligodendrocyte_1 | 923 | 817 | 106 | 88.52 | 135 | 16.52 | 266 | 32.56 | 209 | 25.58 | 207 | 25.34 |
| Oligodendrocyte_2 | 912 | 775 | 137 | 84.98 | 218 | 28.13 | 183 | 23.61 | 201 | 25.94 | 173 | 22.32 |
| Astro_Micro | 881 | 730 | 151 | 82.86 | 129 | 17.67 | 224 | 30.68 | 164 | 22.47 | 213 | 29.18 |
| GABA_nueron | 728 | 511 | 217 | 70.19 | 111 | 21.72 | 116 | 22.70 | 139 | 27.20 | 145 | 28.38 |
| Astrocyte_4 | 605 | 487 | 118 | 80.50 | 109 | 22.38 | 122 | 25.05 | 126 | 25.87 | 130 | 26.69 |
| Astrocyte_5 | 564 | 470 | 94 | 83.33 | 82 | 17.45 | 127 | 27.02 | 147 | 31.28 | 114 | 24.26 |
| Endothelial | 516 | 359 | 157 | 69.57 | 84 | 23.40 | 104 | 28.97 | 80 | 22.28 | 91 | 25.35 |
| Astro_Endo | 506 | 453 | 53 | 89.53 | 81 | 17.88 | 128 | 28.26 | 100 | 22.08 | 144 | 31.79 |
| Tanycyte | 356 | 256 | 100 | 71.91 | 56 | 21.88 | 47 | 18.36 | 85 | 33.20 | 68 | 26.56 |
| Astrocyte_6 | 261 | 216 | 45 | 82.76 | 46 | 21.30 | 54 | 25.00 | 63 | 29.17 | 53 | 24.54 |
| Ependymocyte | 253 | 201 | 52 | 79.45 | 42 | 20.90 | 32 | 15.92 | 59 | 29.35 | 68 | 33.83 |
| Vascular_Leptomeningeal | 239 | 177 | 62 | 74.06 | 44 | 24.86 | 39 | 22.03 | 35 | 19.77 | 59 | 33.33 |
| Microglia_2 | 80 | 64 | 16 | 80.00 | 8 | 12.50 | 14 | 21.88 | 14 | 21.88 | 28 | 43.75 |
| T_cell | 76 | 64 | 12 | 84.21 | 10 | 15.63 | 13 | 20.31 | 21 | 32.81 | 20 | 31.25 |
|  | 20923 | 17396 | 3527 | 80.47 | 2835 | 19.07 | 4473 | 24.27 | 5098 | 27.18 | 4990 | 29.48 |

Analysis of the total cell population revealed that 83.2% (17 396 cells) carried the multiplexing sample tags (Figure 4b and figure S4a). Demultiplexing of this population uncovered 2 835 cells for WT hypothalamus (19.07% of total tagged cells), 4 473 cells for 3xTg hypothalamus (24.27% of total tagged cells), 5 098 cells for WT LVW (27.18% of total tagged cells), and 4 990 cells for 3xTg LVW (29.48% of total tagged cells) (Figure 4 c-d and Table 3), thereby confirming effectiveness of the multiplexing strategy. We then calculated the tagging efficiency across cell types (Figure 4e and Table 3) and found that this ranged from a minimum of 69% (endothelial) to a maximum of 97.5%. On average, the tagging efficiency was 80.47% across all cell types, with the highest efficiency of tagging observed for astrocytes subcluster 1 (97.5%), oligodendrocytes subcluster 1 (88.52%), and the Astro_Endo mixed cluster (89.53%) (Figure 4e). Sample-specific differences in cell type representation were identifiable, such as astrocyte subcluster 1 being increased by 174.24% in the 3xTg hypothalamus versus WT hypothalamus, and astrocyte subcluster 3 being increased by 72.27% in the 3xTg LVW versus WT LVW (Figure S4b). Region-specific differences in WT versus 3xTg gene expression were likewise detectable at the single-cell level (Figure 4f). For example, astrocyte subclusters 1 and 3 had 2-fold more downregulated genes in the 3xTg LVW than in the hypothalamus, while astrocyte subcluster 4 had 6-fold more downregulated genes in the 3xTg hypothalamus than in the LVW (Figure 4f). In both regions, changes in microglial gene expression at this early 4-month-old timepoint were specifically restricted to microglia subcluster 2, a subtype representing only 2.9% of the total microglial population.

Collectively, these data demonstrate that the Thy1.2-based multiplexing strategy permits efficient sample tagging and recovery of all mouse brain cell types for transcriptomic analyses.

## DISCUSSION

This study reports the design and testing of the first antibody-based strategy for recovering all major adult mouse brain cell types following transcriptome sequencing of multiplexed single-cell RNA sequencing samples. The protocol was established for use with the BD Rhapsody microwell-based single-cell capture system and the BD FLEX-Single-cell Multiplexing Kit, the latter enabling use of any PE-labelled primary antibody for multiplexing of samples. Among the major benefits of the protocol are the reduced experimental costs with an optimized pooling strategy (one run with multiple samples at the same time), the increased data yield per experiment (all brain cell populations can be recovered from every experiment for transcriptomic analysis), the elimination of sequencing batch effects (a single sample contains multiple or all experimental groups), and the relative simplicity and reliability of the two-step multiplexing procedure. When combined with the benefits of the BD Rhapsody system, which is gentle and resistant to blockage (gravity-rather than microfluidic-based), has an integrated scanner for precise quantification of the actual number of single cells obtained (enabling exact calculation of the required sequencing parameters), and a relatively simple and user-friendly interface, this protocol represents a cost-effective and reliable strategy for transcriptomic studies of the adult mouse brain. We highlight three specific features of the present study.

First, we used the BD Rhapsody scanner to optimize a number of parameters of our adult brain cell dissociation protocol. Taking into account recent publications, the parameters tested included the cell dissociation temperature, the use of transcription inhibitors, the number and time of washing steps, and cell isolation and purification steps. The final cell dissociation protocol thus decreases processing time and cell stressors as much as possible and resulted in consistently high cell viability and concentration from the multiple microdissected regions of the adult mouse brain.

Second, we identified anti-Thy1.2 as a primary antibody that tags the majority of all major cell types in the adult mouse brain. By flow cytometry, more than 95% of all cells dissociated from the adult mouse brain using the above dissociation protocol were labelled by anti-Thy1.2. In comparison, less than 30% of brain cells were labelled by ACSA-2, Notch-1, CD140a, NRP-1, MHC 1, CD24 or CD13. The labelling of most dissociated brain cells by anti-Thy1.2, also known as CD90.2, was unexpected as it is typically considered a marker of neurons in the adult brain. When PE-conjugated anti-Thy1.2 was used in conjunction with the BD FLEX-Single-cell Multiplexing Kit, the majority of neurons, astrocytes, oligodendrocytes, ependymocytes, tanycytes, microglia, leptomeningeal and endothelial cells were all recoverable for transcriptomic analysis following demultiplexing. The same approach used for identifying anti-Thy1.2 can be used to uncover new multiplexing targets for other organs and tissues.

Third, we performed a proof-of-concept multiplexing experiment using two strains of mice (3xTg-AD and its WT strain control) and two brain regions (hypothalamus and LVW), yielding a single sample containing four experimental groups for sequencing. Over 83% of the combined cells from these groups carried their respective sample tags and could therefore by successfully demultiplexed. Analysis of the demultiplexed samples confirmed that sample-specific differences in cell type representation could be detected, as well as region-specific and cell-specific transcriptomic differences. Thus, the multiplexing protocol described here enabled transcriptomic analysis of all major cell types from four experimental groups in a single run.

Interestingly, BD has recently updated their system, introducing a new cartridge with 8 individual lanes, the HT (high throughput) single-cell system. This new cartridge allows upscaling of experiments to recover up to 55 000 cells per lane (440 000 cells per cartridge). Application of the multiplexing strategy described here to the newest version of the Rhapsody cartridge will allow for even higher recovery of transcriptomic information.

---

**Supplementary figure 1:**
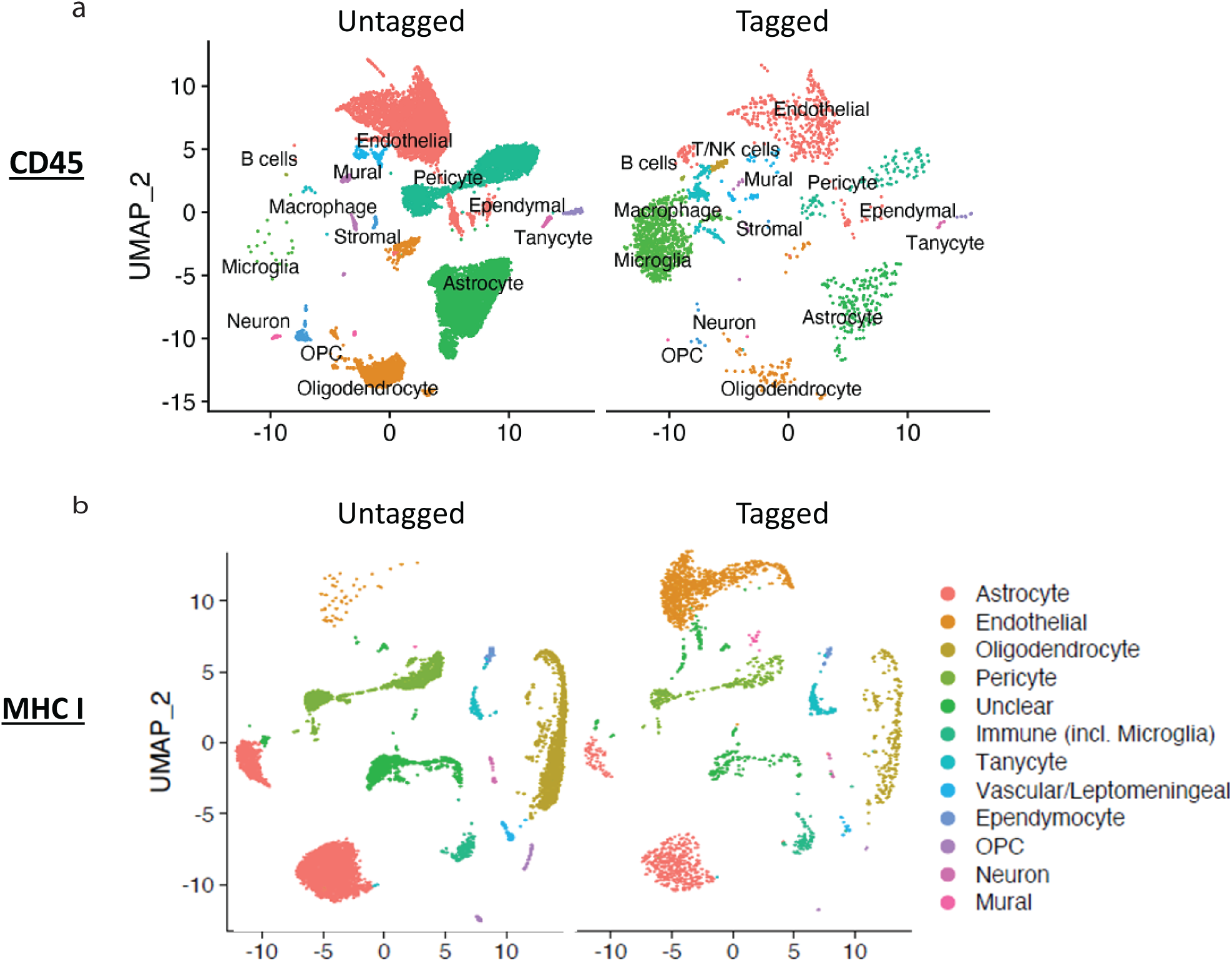
a) CD45 tagging efficiency in adult mouse brain hippocampus. In the tagged clusters, the only reliably tagged group is the microglia cluster. b) MHC-I tagging efficiency in adult mouse brain hypothalamus and lateral ventricle wall. In the tagged clusters, only endothelial cells are consistently identified with the multiplexed groups.

**Supplementary figure 2:**
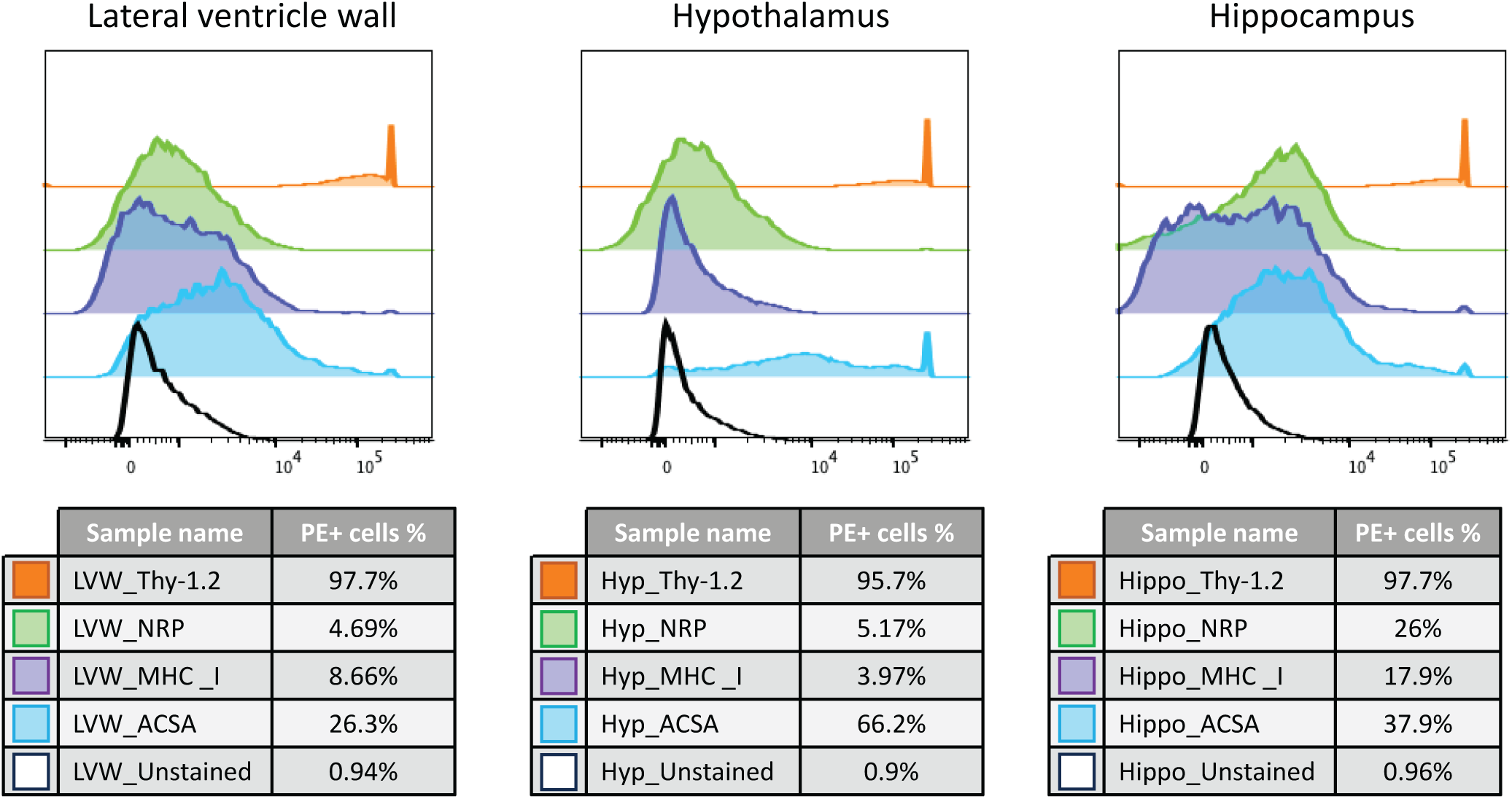
Modal option scales of all channels as a percentage of the maximum count in 3 brain regions: LVW, hypothalamus, and hippocampus.

**Supplementary figure 3:**
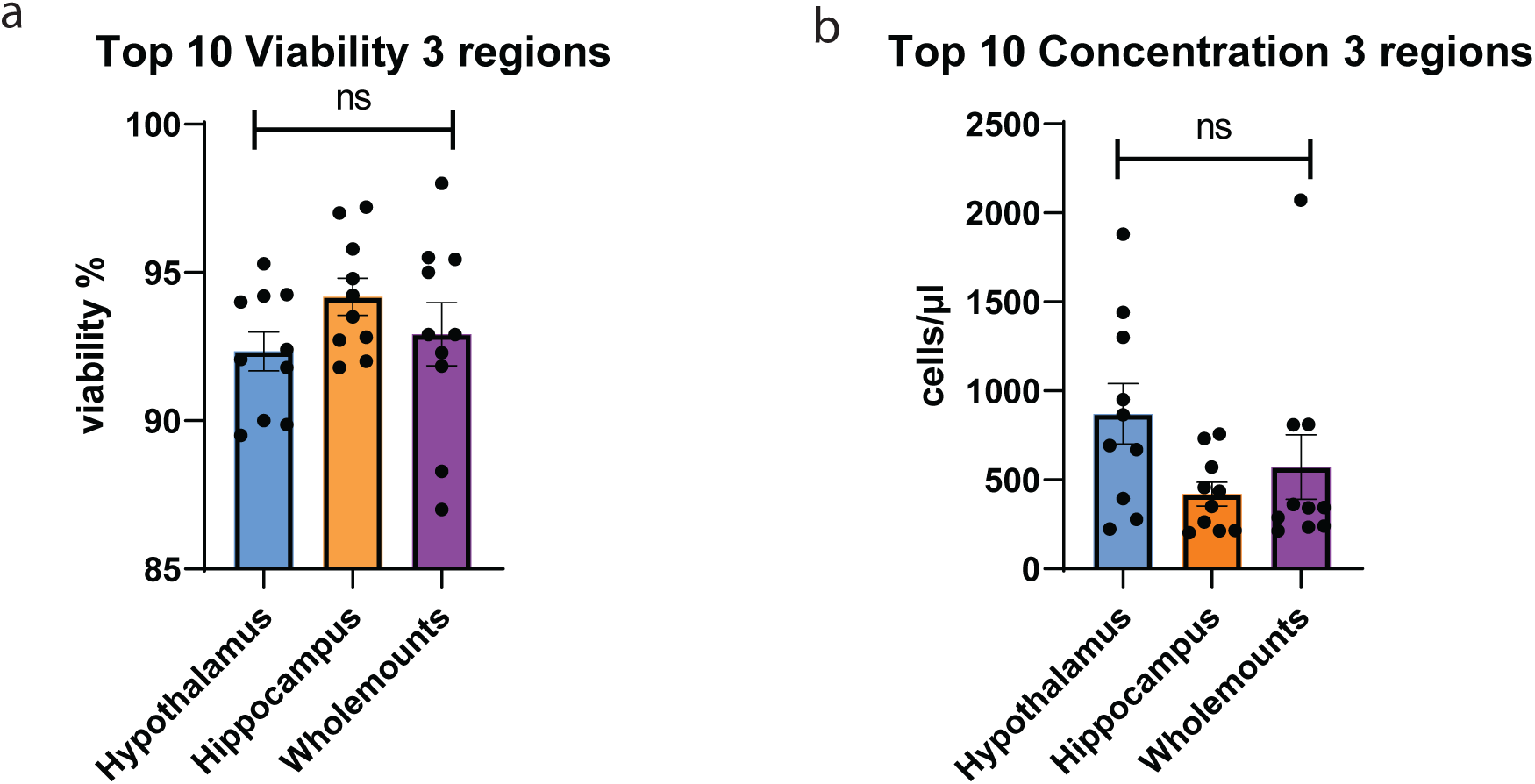
a) Viability percentage among the three brain regions. b) Concentration of cells among the three brain regions.

**Supplementary figure 4:**
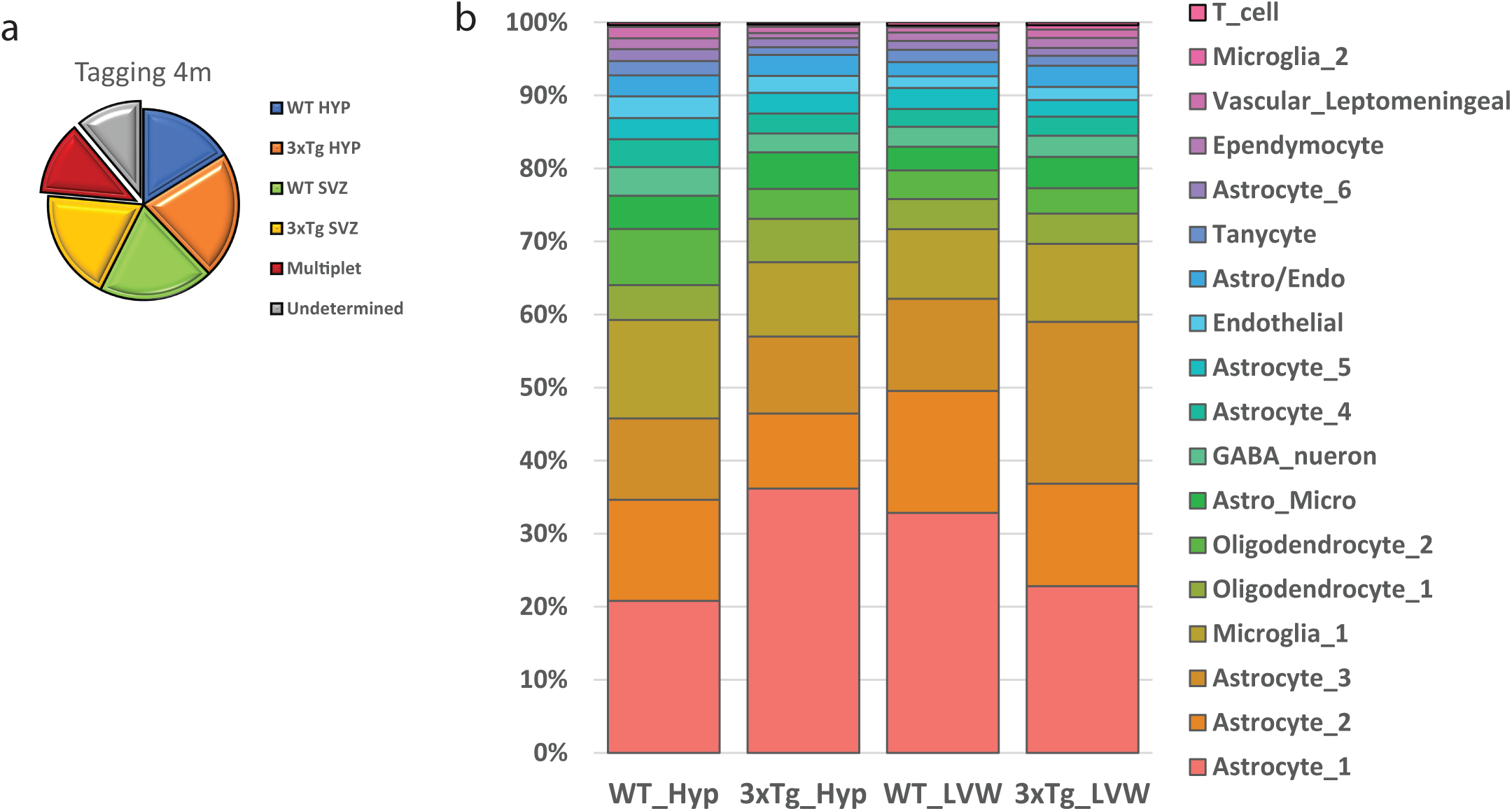
a) Pie chart representing numbers and percentages of cells per group after alignment in Seven Bridges. b) Relative distribution of cell types within the four experimental groups.

**Supplementary table 1:** Original table used to screen the possible candidates for the two-step multiplexing labeling. Some antibodies did not have a PE-conjugated version, therefore they had to be excluded.

| Cell Type | Ab already tested | Ab to test | Clone | Catalogue number |
| --- | --- | --- | --- | --- |
| Astrocyte |  | GLT1 | A18128E | Biolegend - Only Purified format available |
|  |  | AQP4 |  |  |
| Endothelial | <u>MHC I</u> |  | M1/42 | 566776 |
|  |  | PECAM-1/CD31 | MEC 13.3 | 561073 and 553373 |
|  |  | ZO-1/Claudin5 |  |  |
| Ependymocyte |  | CD24 (567383) | 30-F1 | 567383 |
|  |  | CD133 | 315-2C11 | Biolegend (141203 and 141204) |
| Immune (incl. Microglia) | <u>CD45</u> |  | 30-F11 | 553081 |
|  | <u>MHC I</u> |  | M1/42 | 566776 |
| Mural/smooth muscle cells |  |  |  |  |
| Neuroblast |  | CD56 or NCAM1 | 809220 | we have, not in PE |
|  |  | DLX2 |  |  |
| Neuron |  | THY-1 (Thy-1.2) | 53-2.1 | 553005 |
|  |  | THY-1 (Thy-1.1) (554898) | OX-7 | 551401 and 554898 |
| NIP (intermediate progenitors) |  | CD56 or NCAM1 | 809220 | we have, not in PE |
| Oligodendrocyte |  | A2B5 | 105/A2B5 | Purified and Alexa Flour 647 |
|  |  | GALC-MOG |  |  |
|  |  | CNP (is this CNPase? if so - > ) | SMI 91 | Biolegend - Purified and Alexa Fluor 647 |
|  |  | nogoA |  |  |
| OPC (oligodendro. Progenitor cells) |  |  |  |  |
| Pericyte |  | NG2 | MEL62 | Biolegend - Only Purified format available |
|  |  | CD13 | R3-242 | 558745 |
|  |  | PDGFRA | APA5 | 562776 |
|  |  | PECAM-1(CD31) | MEC 13.3 | 561073 and 553373 |
| Tanycyte | <u>MHC I</u> |  | M1/42 | 566776 |
| Vascular/Leptomeningeal (meningeal) |  | NRP-1 | #3E12 | Biolegend - 145203 and 145204 |

## MATERIALS AND METHODS

### KEY RESOURCES TABLE

| REAGENT or RESOURCE | SOURCE | IDENTIFIER |
| --- | --- | --- |
| **Chemicals, Peptides, and Recombinant Proteins** |  |  |
| Narketan (Ketamine 100 mg/ml) | Bimeda-MTC | DIN 02374994 |
| Rompure (Xylazine 20 mg/ml) | Bayer Healthcare | DIN 02169592 |
| Phosphate buffered saline (PBS) | Wisent | Cat#311-010-CL |
| Hank's Balanced Salt Solution (HBSS) | Wisent | Cat#311-512-CL |
| Papain | Worthington | Cat#LS003126 |
| DNase I | Sigma | Cat#DN25 |
| Trypsin Inhibitor type II-O | Sigma | Cat#T9253 |
| DMEM + GlutaMAX | Invitrogen | Cat#10 567 014 |
| F12 + GlutaMAX | Invitrogen | Cat#31 765 035 |
| Hybrimax red blood cell lysis buffer | Sigma | Cat#R7757 |
| CalceinAM | Life | Cat#C1430 |
| DRAQ7 | BD Biosciences | Cat#564904 |
| Stain buffer (FBS) | BD Bioscience | Cat#554656 |
| EDTA 0,53 mM | Wisent | Cat#325-060-EL |
| L-cysteine | Bioshop | Cat#CYS342.10 |
| Actinomycin D | Sigma-Aldrich | Cat#A1410 |
| Triptolide | Sigma-Aldrich | Cat#T3652 |
| Anisomycin | Sigma-Aldrich | Cat#A9789 |
| **Critical Commercial Assays** |  |  |
| Dead-cell removal kit | Miltenyi Biotech | Cat#130-090-101 |
| **Deposited Data** |  |  |
| Single-cell RNAseq experiment | GEO deposition |  |
| **Experimental Models: Organisms/Strains** |  |  |
| Mouse: B6;129-Tg(APP<sup>Swe</sup>,tauP301L)1Lfa *Psen1*<sup>tm1Mpm</sup>/Mmjax | Jackson Laboratories | MMRRC Stock No: 34830-JAX |
| Mouse: B6129SF2/J | Jackson Laboratories | Stock No: 101045 |
| **Other** |  |  |
| 40 µm strainers | Fisher | Cat#22363547 |
| 0.2 µm syringe filters | Sarstedt | Cat#83.1826.001 |
| Plastic pipettes | Fisher | Cat#13 711 48 |
| Hemocytometer | INCYTO | Cat#DHC-N01 |
| Centrifuge | Eppendorf | Cat#2231010060 |
| Isotemp 210 Digital Water Bath | Fisher | Cat#15-462-10Q |
| MACs holder | Miltenyi Biotech | Cat#130-042-303 |
| MACs magnet | Miltenyi Biotech | Cat#130-042-108 |
| MS columns | Miltenyi Biotech | Cat#130-042-201 |
| 50 ml tubes | Sarstedt | Cat#62.547.205 |
| 15 ml tubes | Sarstedt | Cat#62.554.205 |
| 10 ml pipettes | VWR | Cat#CA53300-523 |
| 1.5 ml Eppendorf tube | Fisher | Cat#05 408 129 |
| Filtered and sterile tips 10 µl | VWR | Cat#76322-132 |
| Filtered and sterile tips 20 µl | VWR | Cat#76322-134 |
| Filtered and sterile tips 300 µl | VWR | Cat#10017-088 |
| Filtered and sterile tips 1000 µl | VWR | Cat#10017-090 |

### Resource availability

#### Lead contact

Further information and requests for resources and reagents should be directed to and will be fulfilled by the lead contact, Karl Fernandes.

#### Experimental model and study participant details

##### Mice

Experiments were approved by the Institutional Animal Care Committee of the Pavillon De Recherche Appliquée Sur Le Cancer (Institut de recherche sur le cancer de l’Université de Sherbrooke) following the Canadian Council of Animal Care guidelines. For this experiment we used 3xTg mice (B6;129-Tg(APPSwe,tauP301L)1Lfa Psen1tm1Mpm/Mmjax, Jackson Laboratory MMRRC stock #: 034830) and their WT strain controls (B6129SF2/J, Jackson Laboratory stock #: 101045). Female mice were used for all experiments and sex and age matched.

#### Method details

##### Dissection and microdissection

Mice received a lethal dose of ketamine (Bimeda-MTC)/ xylazine (Bayer Healthcare) and were then perfused transcardially with phosphate-buffered saline (PBS) followed by rapid dissection of the brain, and microdissection of hypothalamus, lateral ventricle wall and/or hippocampus into ice cold HBSS.

##### Single cell dissociation

Preparation of solutions - TIMING (1 hour)

1. Set water bath temperature to 37°C.
2. Set centrifuge (suitable for 15 ml tubes and with option to control braking speed) temperature to 4°C. We maintain the acceleration at level 9 and decrease the break to level 2 (this level depends on the model of centrifuge).
3. Prepare L-Cysteine (100 mM).

a. Add 0.17563 g L-cysteine to 10 ml H2O.
b. Filter on 0.2 µm syringe filter.
c. Store at 4°C (for up to 3 weeks).
4. Prepare Papain activation solution.

a. 10 ml of EDTA 0.53 mM.
b. 100 µl of L-Cysteine 100 mM.
c. Store at 4°C (for up to 3 weeks).
5. Prepare DNase I.

a. NaCl (0.15 M).

1. 0.438 g in 50 ml water.
2. Filter on 0.2 µm syringe filter.
3. Store at 4°C.
b. DNase I (2000 U/ml).

i. Dissolve 15 000 U of DNase I in 500 µl of 0.15 M NaCl.
ii. Stir frequently and keep on ice for 30 minutes.
iii. Transfer to a 15 ml tube and add 7ml of 0.15 M NaCl.
iv. Shake for an hour at 4°C.
v. Filter on 0.2 µm syringe filter.
vi. Aliquot 105 µl/vial.
vii. Store stock solution at −20°C.
6. Prepare Ovomucoid– Trypsin inhibitor (4 mg/ml).

a. Add 0.2 g Ovomucoid– Trypsin inhibitor type II-O to 50 ml HBSS.
b. Shake for an hour at 4°C.
c. Filter on 0.2 µm syringe filter.
d. Aliquote 1.2 ml/vial.
e. Store stock at −20°C.
7. Prepare Papain stock solution.

a. Dilute 1 bottle (1 ml) in 6 ml of HBSS.
b. Store at 4C and protected from light.
8. Prepare Papain/DNase I solution (use within 2 hours): 500 µl of Papain/DNase I per hypothalamus, hippocampus or wholemount (prepare volume according to the number of dissections you will have).

a. Add 15 µl Papain per 1 ml Papain activation solution.
b. Add 100 µl DNase I per 1 ml Papain activation solution.
c. Aliquot each 1 ml in a 15 ml tube.
d. Place all 15 ml tubes in the water bath at 37°C for activation.
e. Note: it is important to activate the enzyme at 37°C for at least 30 minutes before the dissociation on ice to ensure maximum efficiency. At 4°C the cells will show less *ex vivo* activation signature, while the activated enzyme will still be able to function.
9. Prepare solutions with transcriptomic inhibitors. Note: Inhibitors can be avoided when practicing the dissociation protocol. **Actinomycin D** (Sigma-Aldrich, cat. no. A1410) was reconstituted in dimethylsulfoxide (DMSO) at stock concentration of 5 mg/ml and aliquoted and stored at −20°C, protected from light.

- Add 1 ml of DMSO to the original vial and dissolve the crystals.
- Aliquot 50 µl in 20 Eppendorf tubes before storing them at −20°C. **Triptolide** (Sigma-Aldrich, cat. no. T3652) in dimethylsulfoxide (DMSO) at stock concentration of 10 mM was aliquoted and stored at −20 °C, protected from light.
- Add 277.47 µl of DMSO to the original vial and dissolve the powder.
- Aliquot 50 µl in 6 Eppendorf tubes before storing them at −20°C. **Anisomycin** (Sigma-Aldrich, cat. no. A9789) was reconstituted in dimethylsulfoxide (DMSO) at stock concentration of 10 mg/ml and aliquoted and stored at +4 °C, protected from light.
- Add 500 µl of DMSO to the original vial and dissolve the powder.
- Aliquot 50 µl in 10 Eppendorf tubes before storing them at +4°C. Perfusion buffer (perfusion buffer + inhibitors):
- 20 ml per mouse plus 10 ml per channel of perfusion (example for 2 mice).

1. Add 60 ml of cold HBSS.
2. Add 60 µl of Actomicin D (5 μg/ml, 1:1 000 from stock).
3. Add 60 µl of Triptolide (10 μM, 1:1 000 from stock). Dissection buffer (dissection buffer + inhibitors): *Prepare fresh before each experiment
- 1 ml per tube with 2 HPP. (Multiply the volume according to the tubes needed and do a stock solution)
- Add inhibitors immediately before beginning perfusion. Keep on ice protected from light.

1. Add 1 ml of cold HBSS.
2. Add 1 µl of Actomicin D (5 μg/ml, 1:1 000 from stock).
3. Add 1 µl of Triptolide (10 μM, 1:1 000 from stock).
4. Add 2.7125 µl of Anisomycin (27.1 μg/ml, 1:368.5 from stock). Digestion buffer (digestion buffer + inhibitors): *Prepare fresh before each experiment
- 1 ml per tube with 2 HPP (Multiply the volume according to the tubes needed and do a stock solution)
- Add inhibitors immediately before beginning perfusion. Keep on ice protected from light.

1. Add 1 ml of cold HBSS.
2. Add 1 µl of Actomicin D (5 μg/ml, 1:1 000 from stock).
3. Add 1 µl of Triptolide (10 μM, 1:1 000 from stock).
4. Add 2.7125 µl of Anisomycin (27.1 μg/ml, 1:368.5 from stock).
10. Prepare tools for brain dissection.

### Step-by-Step Method Details

#### Brain dissection

TIMING: depending on dissection method, approximately 5-7 minutes per mouse for hippocampal and hypothalamus dissection when a team of 3 people is involved.

1. Inject a lethal dose of ketamine (347 mg/kg) /xylazine (33 mg/kg) mixture.
2. When the mouse shows no reflexes, begin transcardial perfusion with 20 ml of phosphate-buffered saline (PBS) or HBSS + inhibitors.
3. Immediately decapitate and dissect the brain into a dish of HBSS (on ice).
4. When performing microdissections, place the brain in a dish containing HBSS (on ice) and transfer to Eppendorf tube on ice.
5. Pool 2 hypothalami, 2 hippocampi or 2 lateral ventricle walls into 1.5 ml Eppendorf tube containing 1ml of HBSS (keep on ice).

**Critical:** To achieve maximum viability, dissections should be done as quickly as possible, and cells should be kept on ice when indicated. If many mice are required for your study, we suggest you work as a team with one person injecting the mice and perfusing multiple mice (with an appropriate delay to allow for immediate dissection), one person decapitating and removing the whole brain, and one person performing the microdissections. Working this way, we have dissected one hippocampus and the hypothalamus from 8 mice in approximately 1 hour. For our experiments we needed to dissociate 8 tubes (containing 2 microdissections) simultaneously, so we were two people dissociating 4 tubes each. We do not recommend dissociating more than 4 tubes at a time as this will increase the total time of the experiment and possibly decrease viability.

**Critical:** We recommend dissociating between 20-35 mg of tissue per 15 ml tube containing 1 ml of Papain/DNase I. In our experience we observed no differences in viability using 2 hippocampi (35 mg), 2 lateral ventricle walls (20 mg) or 2 hypothalami (35 mg) of tissue per 15 ml tube. It is possible that using more or less tissue will be acceptable, but we have not tested this.

### Cell Dissociation

TIMING (approximately 1.5 hours but dependent on number of samples). Once the microdissections are all collected in cold HBSS:

1. Open all Pasteur plastic pipets and wet it with Papain/DNase solution.
2. Transfer microdissections into 15 ml tubes containing 1 ml Papain/DNase I with a plastic pipette.
3. Triturate with 20/25 up and down using the plastic pipette. Note: Start by aspirating a little bit of liquid and tap on the tissue until it can be easily aspirated inside the plastic pipet. Only then, start counting 20/25x reaching only half of the pipet length. It is normal to see pieces of tissue at this step.
4. Incubate 10 mins ON ice (4°C).
5. Using P1000 (set at 800 µl) and low binding tips triturate with 20/25 up and down.
6. Incubate 10 mins on ice (4°C). While the incubation takes place:
7. Prepare Base Media (BM) (in a sterile container like a cell culture flask)

- 3 parts DMEM + 1 part of F12
- For 1 tube prepare a total volume of 37.5 ml.
- Multiply the quantity depending on the number of tubes per user: For 1 tube: 28.125 ml of DMEM + 9.375 ml of F12 For 4 tubes: 112.5 ml of DMEM + 37.5 of F12.
8. Prepare the 40 µm strainer on top of a 50 ml tube.
9. Take the tube in incubation (step 6) and do 20/25 up and down with a P1000 set at 800 µl.
10. Add 1 ml of Ovomucoid x Tube (1:1 ratio).
11. Do 20/25 up and down with a P1000 set at 800 µl.
12. Do 20/25 up and down with a P200 set at 200 µl.
13. Add BM up to 9 ml and mix the closed tube by flipping it upside down a couple of times.
14. Wet the 40 µm strainer with 3 ml of BM before passing the cell suspension through it (prepared at step 11.)
15. Remove with pipetman the 3 ml of BM used to wet the first strainer and use it to wet the next strainer in case you have multiple tubes. Dispose it in the liquid waste container once all strainers are wet.
16. Pour the 9 ml of cell suspension into the 50 ml tube through the stainer.

- Make sure the entire sample volume is poured through the strainer before disposing the 15 ml tube.
17. Rinse the strainer with 5 ml BM.
18. Pass the filtered sample to a new 15 ml tube. (Do it by hand but decant by swirling to transfer all cells).
19. Centrifuge for 10 minutes at 4°C 1500 rpm/500 g without brakes.
20. During this centrifugation, prepare 1:1 dilution of the red blood cell lysis buffer solution. Per tube prepare the following:

- 2.5ml of BM.
- 2.5 ml of red blood cell lysis buffer. For 4 tubes: 10 ml of BM + 10 ml of red blood cell lysis buffer. Keep on ice.
21. After centrifugation, take the 15 ml tubes.
22. Carefully remove supernatant without disturbing the pellet, and leave 1 ml. Note: Use a 10 ml pipet until 2 ml and use a P1000 to remove 1 ml.
23. Gently resuspend the cells with P1000. Note: Do step 26-27 for each tube before adding the red blood cell lysis buffer to all the samples.
24. Add 5 ml of red blood cell lysis buffer and incubate for 1 minute (6 ml tot). Coordinate with the working partner to put the red blood cell lysis buffer at the same time. Note: In case there are 4 tubes, pipet 10 ml to do 2 samples and repeat again for the 2 other samples. Close the tube and invert to mix well before starting the timer 1 min.
25. Add BM up to 14 ml.
26. Centrifuge for 10 minutes at 4°C 1500 rpm/500 g without brakes.
27. From the centrifuged tube, carefully remove supernatant and leave 1 ml. Note: Use a 10ml pipet until 2 ml and use a P1000 to remove 1 ml.
28. Gently resuspend the cells with P1000. Note: Do steps 31-32 for each tube before adding the BM to all the samples.
29. Add BM up to 6 ml.

Pooling step

1. Prepare the 40 µm strainer on top of a 50 ml tube.
2. Wet strainer with 3 ml of BM before passing the cell suspension through it.
3. Remove with pipetman the 3 ml of BM used to wet the strainer and use it to wet the next strainer in case you have multiple tubes. Dispose in the liquid waste container.
4. Pass the cell suspension through it to remove cell clumps. Pool 2 tubes together. Note: Make sure the entire sample volume is poured through the strainer before disposing the 15 ml tube.
5. Rinse strainer with 3 ml BM. Note: No piece of tissue should be visible at this step
6. Transfer in a new labeled 15 ml tube and mark where the 0.5 ml line is.
7. Centrifuge for 10 minutes at 4°C 1500 rpm/500 g without brakes.
8. Aspirate supernatant as much as possible. Note: Use a 10ml pipet until 2 ml and use a P1000 to remove as much as possible
9. Add 100 µl of Dead cell removal beads and resuspend. (find colorful purple box in 4°C fridge in front of the sink)
10. 10. Incubate 15 mins on ice (4°C). During this incubation:
11. Prepare 1x Binding Buffer (keep on ice): For 1 column: 4 ml total

- 3.8 ml H2O (use sterile commercial water).
- 220 µl of 20x Binding buffer. For 4 columns: 15.2 ml of H2O + 880 µl of 20x Binding buffer.
12. 5 minutes before end of incubation, place columns on magnet. Note: Make sure the lines from the column are looking towards the user to assure the column is placed straight in the holder.
13. Rinse columns with 500 µl 1x binding buffer (x2)
14. Empty the 15 ml tubes of Binding Buffer, after rinsing the columns. At the end of the 15 minutes incubation:
15. Add 1x binding buffer to cells up to 500 µl (sample tube).
16. Place a new labeled 15 ml cell collection tubes under columns.
17. Set your P1000 at 600 µl before resuspending the cells with 3 to 5 gentle ups and downs.
18. Apply the entire volume of cell suspension on corresponding column.
19. Let the live cells pass through into the 15 ml collection tubes by gravitational force.
20. Rinse column 4x with 500 µl 1x binding buffer.
21. Centrifuge cells 10 minutes at 4°C 1500 rpm/500 g without brakes. Note: it is possible to quantify viability and concentration of your sample at this stage using Calcein Am and Draq7 staining. We recommend training up to this point several times before handling the real samples to ensure quality and quantity of the single cell suspension.

**Cell staining using Multiplexing set – Flex SMK anti-PE**

1. Remove all supernatant using P1000 (leave 2 mm on top). Note: aspirate as much as possible, but do not take risks when the pellet is very small.
2. Resuspend cells up to 190 µl FBS buffer (BD) for Multiplexing. Note: make sure you keep into consideration the leftover volume on top of the pellet.
3. Pipette 190 µl of cell suspension into 5 ml polystyrene low binding tube. Note: we achieved similar results using the 15 ml tubes.
4. Add 10 µl of anti-Thy1.2-PE antibody from stock tube (following instruction from manufacturer). Note: All tubes will have the same Ab at this point.
5. Do gentle up and down to mix.
6. Incubate at 4°C 20 min in the dark.
7. Add 1 ml of cold FBS buffer (BD) to the 5 ml polystyrene low binding Tube containing the cell suspension and resuspend the pellet.
8. Add an additional 1 ml of FBS buffer (BD) for a total of 2.2 ml.
9. Mix well by inversion.
10. Centrifuge each tube 5 minutes at 4°C 1500 rpm/500 g without brakes.
11. Use P1000 to remove supernatant without disturbing the pellet.
12. Add 1 ml of cold FBS buffer (BD) to the 5 ml polystyrene low binding Tube containing the cell suspension and resuspend the pellet.
13. Add an additional 1 ml of FBS buffer (BD) for a total of 2 ml.
14. Use P1000 to remove supernatant as much as possible.
15. Resuspend up to 180 µl FBS. Note: make sure you keep into consideration the leftover volume on top of the pellet.
16. Add 20 µl of SMK secondary antibody anti-PE with sample tag (following instruction from manufacturer).
17. Mark which sample tag goes with which sample.

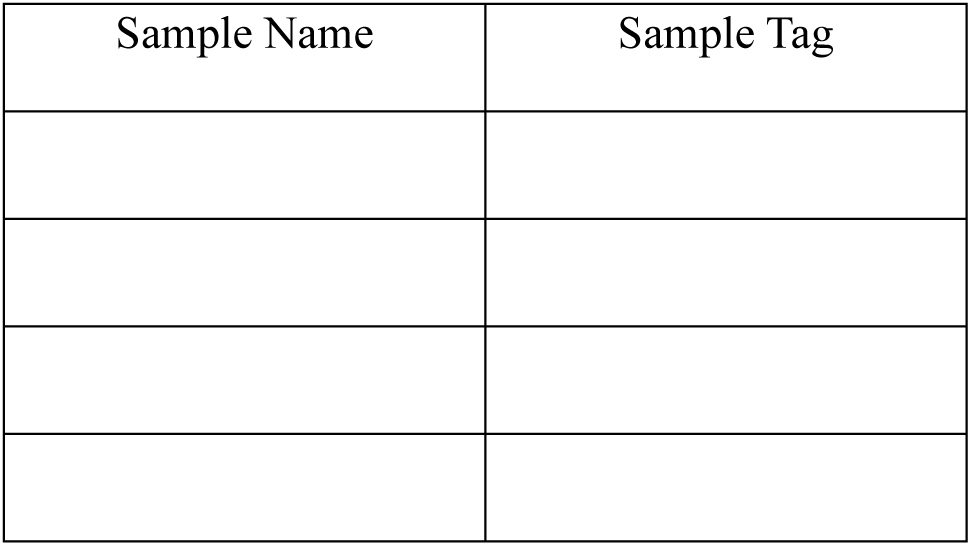
18. Mix with ups and downs.
19. Incubate at 4°C 20 min in the dark.
20. Add 1 ml of cold FBS buffer (BD) to the 5 ml polystyrene low binding Tube containing the cell suspension and resuspend the pellet.
21. Add an additional 1 ml of FBS buffer (BD) for a total of 2.2 ml.
22. Mix well by inversion.
23. Centrifuge each tube 5 minutes at 4°C 1500 rpm/500 g without brakes.
24. Use P1000 to remove supernatant without disturbing the pellet.
25. Add 1 ml of cold FBS buffer (BD) to the 5 ml polystyrene low binding Tube containing the cell suspension and resuspend the pellet.
26. Add an additional 1 ml of FBS buffer (BD) for a total of 2 ml.
27. Mix well by inversion.
28. Centrifuge each tube 5 minutes at 4°C 1500 rpm/500 g without brakes.
29. Use P1000 to remove supernatant without disturbing the pellet.
30. Resuspend cells up to 400 µl cold Sample Buffer from the BD Rhapsody™ Cartridge Reagent Kit with P1000.
31. Add 2 µl of Calcein AM (2mM).
32. Add 2 µl of Draq7 (2mM).
33. Incubate at 37°C for 5 minutes before counting with BD Rhapsody Hemocytometer.

#### Flow cytometry

Eight hippocampi were microdissected from 8-months old female WT mice. Following the single cell dissociation protocol, cells were stained with Calcein AM (494/517 nm, live cells) and Draq7 (647/697 nm, dead cells) in 400 µl of FBS (stain buffer, BD Biosciences) at 37°C for 5 minutes. Cell viability and concentration were quantified on the BD Rhapsody scanner. Cells were then resuspended at a concentration of 50,000 cells per well in 200 µl of FBS. Cells were not fixed to mimic the single cell isolation protocol. Cell surface staining was performed with each PE-conjugated antibody (565/574 nm) added to the single cell suspensions in two different volumes: 1 µl or 3 µl. A list of antibodies used can be found in table 1. The samples were incubated with the antibodies for 20 minutes in the dark at 4°C. Following incubation, the samples were washed twice with FBS. The percentage of stained cells (PE+) was determined by gating on the living cell population, according to granularity and size, and excluding the PE-cells from the unstained sample. Samples were processed using an LSR II cytometer (BD Biosciences), and the analysis of data was carried out utilizing FlowJo v10.5.3 (Tree Star, Ashland, OR, USA).

#### Single cells capture and sequencing

The viability of cells in all samples was between 90–95% (figure 1c). 15,000 cells per sample, were pooled in a single 1.5 ml Eppendorf tube for a total of 60,000 cells. Single-cell capture was carried out using the BD Rhapsody system (BD Biosciences) on a single lane cartridge following manufacturer instructions (BD Rhapsody WTA kit). Subsequently, the mRNA from each cell was secured to the poly-t attached to a barcoded bead. After the beads were retrieved, cDNA was generated using the cDNA kit from BD. Libraries were prepared using the whole transcriptome analysis (WTA) amplification kit (BD Biosciences) following the manufacturer’s recommendations. The quality of both the WTA and Sample tag libraries was assessed using a DNA screentape D5000 HS on a TapeStation 2200 (Agilent Technologies, Santa Clara, CA, USA), and the concentration was measured using a QuBit 3.0 fluorometer (Thermo Fisher Scientific, Canada). Following this, the WTA and Sample Tag libraries were pooled in a ratio of 98.8% to 1.2%.

The libraries were then sequenced with paired-end 100 bp sequencing on a NovaSeq 6000 (Novaseq 6000 S4 reagent kit (200 cycles), Illumina Inc., San Diego, CA, USA) at the Next-Generation sequencing Platform, Genomics Center, CHU de Québec-Université Laval Research Center, Québec City, Canada. The coverage was about 2600M paired-end reads.

#### Bioinformatics

Sequencing FASTQ pair-end files were processed for quality control and aligned with the BD Rhapsody WTA Analysis Pipeline on the SevenBridges application (BD Biosciences) using the RhapRef_Mouse_WTA_2023-02 based on GRCm38 genome and the gencode vM24 transcriptome annotation.

A raw expression matrix with cellular and sample tag barcodes was recovered directly as a Seurat object and processed using standard R package Seurat v4.3.0.1^30^; it contained 35 000 cells with 39 065 mean reads/cell, and 4 866 mean sample tag reads/cell (including all 4 multiplex tags).

Low-quality cells were treated with following filters: number of detected genes (each single cell has a minimum number of expressed genes > 200), low count depth (each single gene expressed at least in 3 cells or more), high fraction of mitochondrial counts (mitochondrial genes with expression in every single cell < 50%). A higher threshold for mitochondrial genes was used in order to better annotate and cluster the samples. Identification of multiple cells (multiplets) was performed by identifying cells by multiple sample tags and subsequently removed. Our final filtered raw count data comprised a final cell count of 20 923 and 30 204 genes.

Log-normalization was performed to standardize gene counts per cell, ensuring comparability. Biologically meaningful genes were then chosen using the feature selection method vst, with a maximum of 2 000 variable genes selected, for subsequent dimensionality reduction. We then applied a linear transformation of data (scaling and centering), which is a prerequisite for subsequent dimensionality reduction that was performed by principal component analysis (PCA). number of principal components (PCs) to consider for the further analysis was determined by either visualizing at least the first 20 most informative principal components (maximum 500 cells each) and by utilizing a re-sampling JackStraw-like test and by visualizing an elbow plot. In our case we chose to go with the first 8 PCs.

We clustered similar expression profile cell population by first performing a k-nearest neighbor (KNN) graph technique and subsequent Louvain algorithm, in our case, we were able to identify 15 clusters. Final visualization was made with the non-linear dimensional reduction technique such as UMAP.

Clusters were annotated by an automated method using two reference-databases (scMCA mouse cell atlas R package v0.2.0.75 and the “Single-cell transcriptomics of 20 mouse organs creates a Tabula Muris” trained model in scibet R package v0.1.076)^31,32^.

Automatic cell type annotation was conducted using scMCA mouse cell atlas R package v0.2.0 and the “Single-cell transcriptomics of 20 mouse organs creates a Tabula Muris” trained model in scibet R package v0.1.0 and was manually curated.

Different condition group demultiplexing was performed by sub-setting Seurat object based on sample names. Identified sample_names and conditions were the 4 experimental groups, WT_Hyp, 3xTg_Hyp, WT_LVW, and 3xTg_LVW and the undetermined group (which was not included in subsequent differential expression analysis). Undetermined cells cannot be confidently assigned to any sample tag group.

Differential expression was made with Seurat function “FindMarkers”, considering a p-adjusted value threshold < 0.05 by comparing the following groups: 3xTg_Hyp vs WT_Hyp and 3xTg_LVW vs WT_LVW.

## Acknowledgements

We thank the Next-Generation Sequencing Platform of the Genomics Center at CHU de Québec-Université Laval Research Center for the single-cell sequencing, the RNomics platform of the Université de Sherbrooke for quality controls on our libraries, and the Bioinformatics platform of the Biochemistry and Functional Genomics department of the University of Sherbrooke. We also want to thank Emily Chomyshyn from BD Bioscience for her technical support in flow cytometry and constant logistical assistance. J.A.L. is supported by Neurology Service of the Faculty of Medicine and Health Sciences (Université de Sherbrooke), I.A. by a scholarship from the FRQS, and N.S. by a MITACS Globalink Research scholarship. M.A.B is a FRQS Junior 1 Fellow. K.J.L.F. is a Tier 1 Canada Research Chair. This work was funded by operating grants to K.J.L.F. from the Canadian Institutes of Health Research (CIHR) and the Natural Sciences and Engineering Research Council (NSERC).

## Author contributions

F.P. developed the concept, conducted experiments, processed data, and co-wrote the manuscript. L.K.H. contributed to the single-cell dissociation and RNA sequencing protocol. J.A.L. and A.A. conducted experiments. M.A. and N.S. conducted bioinformatic data analysis. I.A. and M.B. provided bioinformatics expertise and discussion. K.J.L.F. co-wrote the manuscript and obtained operating funds for the study.

## Notes

### Competing Interest Statement

The authors have declared no competing interest.

